# Gender- and Age-Based Characterization and Comparison of the Murine Primary Peritoneal Mesothelial Cell Proteome

**DOI:** 10.1101/2024.10.09.617441

**Authors:** Zhikun Wang, Yueying Liu, Reihaneh Safavisohi, Marwa Asem, Daniel D. Hu, Mary Sharon Stack, Matthew M. Champion

## Abstract

Organs in the abdominal cavity are covered by a peritoneal membrane, which is comprised of a monolayer of mesothelial cells (MC). Diseases involving the peritoneal membrane include peritonitis, primary cancer (mesothelioma), and metastatic cancers (ovarian, pancreatic, colorectal). These diseases have gender- and/or age-related pathologies; however, the impact of gender and age on the peritoneal MC is not well evaluated. To address this, we identified and characterized gender- and age-related differences in the proteomes of murine primary peritoneal MC. Primary peritoneal MC were isolated from young female (FY) or male (MY) mice (3-6 months) and aged female (FA) or male (MA) mice (20-23 months), lysed, trypsin digested using S-Traps, then subjected to bottom-up proteomics using an LC-Orbitrap mass spectrometer. In each cohort, we identified >1000 protein groups. Proteins were categorized using Gene Ontology and pairwise comparisons between gender and age cohorts were conducted. This study establishes baseline information for studies on peritoneal MC in health and disease at two physiologic age/gender points. Segregation of the data by gender and age could reveal novel factors to specific disease states involving the peritoneum. [This *in vitro* primary cell model has utility for future studies on the interaction between the mesothelium and foreign materials.]

**SUMMARY STATEMENT:** Many diseases initiate from or involve peritoneal mesothelial cells including peritonitis, primary cancer (mesothelioma) and metastatic cancers. Progression of these diseases is influenced by many host factors including gender and age; however, the influence of these factors on the peritoneal mesothelial cell proteome has not been evaluated. This study provides novel information and identifies proteins exclusive to both male and female young and aged cohorts. Given the importance of the peritoneal mesothelial cell in abdominal homeostasis, and the impact of gender and age on disease progression, these data will be key for future studies examining mesothelium in both health and disease.

## 1. INTRODUCTION

A major abdominal organ is the peritoneum, a vast serous membrane that lines the inner walls of the abdominal cavity and the outside of the visceral organs with a continuous surface area of 1-2 m^2^, nearly equal to that of the skin.[1],[2],[3],[4] The peritoneum consists of two compartments separated by a basement membrane: the mesothelium and supporting loose connective tissue.[2],[3],[5] The mesothelium, a monolayer of mesothelial cells (MC), functions to reduce friction between abdominal organs, supports the homeostasis of the peritoneal cavity, and acts as a protective barrier to the collagen I-rich submesothelial matrix[5],[6] and the underlying abdominal organs. As a result, many peritoneal diseases, including peritonitis, primary cancer (mesothelioma), and metastatic cancer, initiate from interactions with the mesothelium [**Figure 1**].

**Figure 1.**
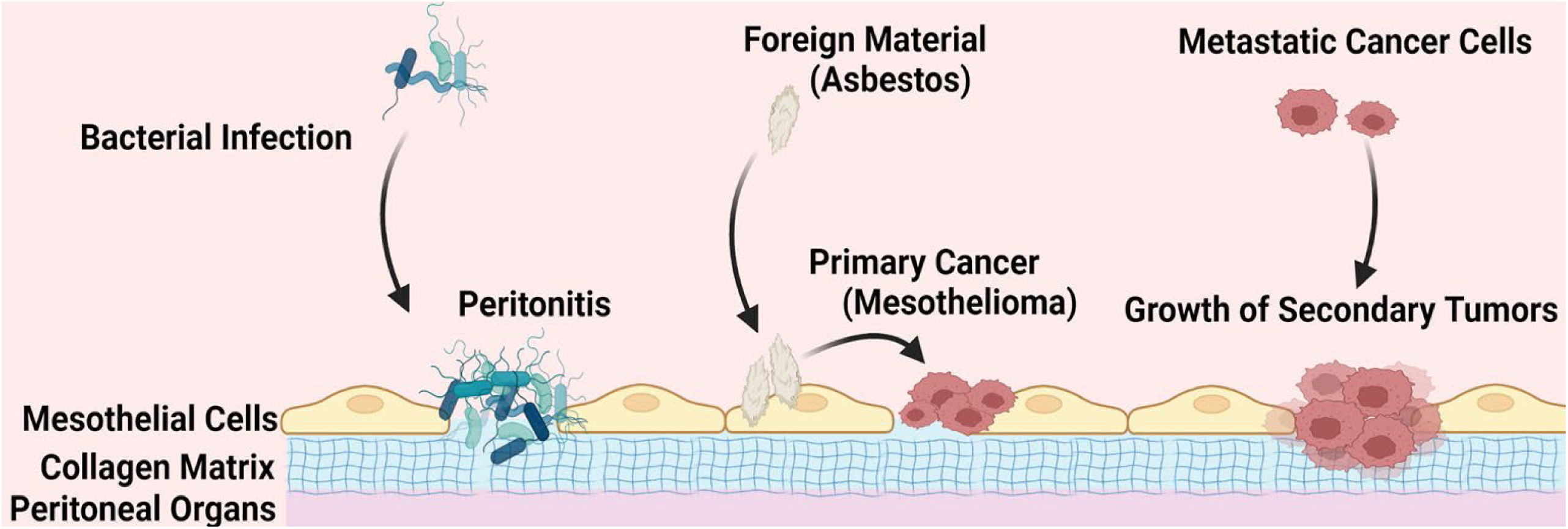
The role of mesothelial cells in peritoneal disease etiology. Several common peritoneal diseases directly involve mesothelial cells. Peritonitis is caused by bacterial infection triggering an inflammatory response on the mesothelium. Mesothelioma is a primary malignancy resulting from direct malignant transformation of mesothelial cells, commonly resulting from asbestos exposure. Secondary malignancies result from adhesion to and metastatic implantation of the mesothelium by disseminating tumor cells, often resulting in mesothelial cell retraction and anchoring of metastases in the sub-mesothelial matrix.

Peritonitis is inflammation of the peritoneum caused by bacterial infection from leakage through the gastrointestinal tract or through the abdominal wall via injury or surgery. It has been shown that advanced age is a major risk factor for the development of post-operative septic complications following abdominal surgery.[7] Sex differences in the inflammatory response have also been described in diverse species.[8] In a study of 745 patients, death due to peritoneal dialysis-associated peritonitis was more common in men compared to women.[9] In murine studies of peritonitis, female mice were shown to recruit fewer classical monocytes and neutrophils.[10]

Mesothelioma is a primary cancer of the mesothelium in which MC become malignant after being exposed to foreign material such as asbestos. It was reported that female mesothelioma patients have improved survival relative to male patients. Moreover, younger patients have better survival than the aged patients, but the age-dependent survival difference is observed only in females.[11] Many other types of cancer, including pancreatic, colorectal, and ovarian cancer produce peritoneal metastasis initiated by adhesion of metastasizing cancer cells to MC. Pancreatic cancer is more common in males than in females. Mortality rates from pancreatic cancer increase with age for both sexes, but overall are higher in aged males.[12] Among 102,998 pancreatic cancer related deaths in Spain, 48,346 (46.94%) were females and 54,652 (53.06%) were males.[12] The age-specific mortality rates per 100,000 increase with age (2.6-72.8 for males, 1.3-60.8 for females) between 40 to 84 years old.[12] Studies on 164,996 colorectal cancer patients in Germany have shown that women have a significantly increased long-term survival rate; however, this advantage disappears after the age of 65.[13] Similar sex-specific differences have also been suggested in studies with patients from Korea, China, and England.[14],[15],[16] Ovarian cancer incidence per 100,000 has also shown significant age-dependent differences. In patients younger than 65 incidence is 9.34, while incidence in older patients is 52.7 in the US.[17],[18],[19] It was also shown that ovarian cancer patients have an overall survival of 47.6 months for women under the age of 65, decreasing to 37.4 months for women ages 65 and older.[17],[18],[19]

While MC are a major factor in many diseases of the peritoneum, little is known regarding the impact of sex and age on MC homeostasis. To probe sex- and age-dependent changes in mesothelial cells, the objective of this study was to isolate murine primary peritoneal mesothelial cells (MPPMC) and perform comparative bottom-up proteomic analysis. The resulting data identify sex- and age-related differences that may regulate MC homeostasis in the healthy peritoneum and influence the initiation and progression of peritoneal diseases.

## 2. MATERIALS AND METHODS

Raw and processed data are available through the MassIVE data exchange and cross-posted to ProteomeExchange. MSV000092134 ftp://MSV000092134@massive.ucsd.edu (For Review) Password: MouseAge2302* https://massive.ucsd.edu/ProteoSAFe/static/massive.jsp

### Materials

Rat tail collagen type I and trypsin were purchased from Corning. Cell culture media compositions include Dulbecco’s Modification of Eagle’s Medium (DMEM, Corning, Midland, MI), Ham’s F12 (Corning), 1% penicillin streptomycin solution (Corning), fetal bovine serum (FBS, Gibco, Carlsbad, CA), epidermal growth factor (EGF, Gibco), hydrocortisone (Corning), GlutaMAX (Gibco), HEPES buffer (Corning), ITS (ITS+1 Liquid Media Supplement, Sigma-Aldrich, St. Louis, MO). Primary antibodies used are vimentin (Sigma-Aldrich, catalog #: V5255) and cytokeratin 18 (catalog #: ab668, Abcam, Cambridge, MA). Anti-mouse IgG peroxidase-conjugated secondary antibodies were from Sigma-Aldrich. Peroxidase detection reagents, SuperSignal West Dura, and Halt™ Protease Inhibitor were obtained from ThermoFisher (Rockford, IL). Ammonium-Chloride-Potassium (ACK) Lysing Buffer was purchased from Gibco. All methods were carried out in accordance with the University of Notre Dame Institutional Review Board (IRB) and Institutional Biosafety Committee (IBC) guidelines and regulations. Experimental protocols were approved by the IBC committee (protocol 23-04-7798, expiration 06/14/26).

### Animals

C57Bl/6 male and female mice were purchased from Jackson Laboratory at 2 age points: mature young (3-6 months old) mice (equivalent to human 20-30 years old) and aged (20-23 months old) mice (equivalent to human 60-67 years old).[19],[20] Cohorts (6 mice per cohort) were defined as Female Young (FY), Female Aged (FA), Male Young (MY) and Male Aged (MA). All animal procedures were carried out according to the regulations of the Institutional Animal Care and Use Committee (IACUC) at the University of Notre Dame. All murine studies were approved by the IACUC and were conducted in accordance with relevant guidelines and regulations of this committee. Additionally, this study is in compliance with the Essential 10 of the ARRIVE guidelines https://arriveguidelines.org/sites/arrive/files/documents/ARRIVE%20Compliance%20Qu estionnaire.pdf. Appropriate details of study design, sample size, outcomes, statistical treatment and methods and experimental animals and procedures are enumerated. No treatments were conducted on the mice; thus, randomization and blinding were not needed/performed for this cohort. No samples were excluded from the study, and all mice were sacrificed equally.

### Murine Primary Peritoneal Mesothelial Cell (MPPMC) Isolation, Culture, and Lysis

Mice were sacrificed via isoflurane overdose in accordance with IACUC guidelines. To isolate primary MCs, immediately after sacrifice, mice were injected intra-peritoneally (i.p.) with 3 mL of 0.125% trypsin. Mice were maintained at 37 for 20 minutes, during which they were gently rotated to mix the peritoneal fluid every 2-3 minutes. After incubation, 6 mL of isolation media (DMEM/F12 1:1, 10% FBS, 1% Pen/Strep) were injected i.p. into each mouse to neutralize the trypsin and the peritoneal fluid was collected. The abdominal cavity was further washed with 3 mL of isolation media and the collected mixture was centrifuged at 180xg at 4. Each cell pellet was re-suspended in 2 mL ACK lysing buffer for 2 minutes on ice to remove blood cells followed by mixing with 6 mL of PBS to terminate the reaction. After centrifugation (180xg, 4 °C), pellets were washed with 6 mL isolation media, followed by another centrifugation (180xg, 4 °C). Cells were re-suspended in culture media (DMEM/F12 1:1, 15% FBS, EGF 10 ng/mL, hydrocortisone 400 ng/mL, 1% Pen/Strep, 1 % L-glutamine, 10 mM HEPES, ITS 1:100) and plated onto tissue culture dishes coated with rat tail collagen type I (10 μg/ml in coating buffer 0.1 M Na_2_CO_3_, pH9.6) or chamber slides as previously described.[21]

In each cohort, 6 mice were sacrificed for MPPMCs and cells from each mouse were plated into individual wells of 24-well collagen-coated plates. Adherent cells were washed with warm PBS at 24 hours after being plated to remove non-adherent and/or dead cells, then cultured for an additional three days prior to changing to serum free medium for 24 hours to remove serum protein contaminants. For proteomic analysis, after washing three times with PBS, cells were lysed by adding 100 μL modified RIPA (mRIPA) buffer (150mM NaCl; 50mM Tris, pH 7.5; 20mM NaF; 10mM Na_2_P_2_O_7_; 5mM EDTA; 1% Triton X-100; 0.1% SDS) containing protease inhibitors to each well for 10 minutes at 4°C. Preliminary analyses showed that the mass balance of these proteins was predominately media and murine blood-associated proteins. Therefore, lysates from each cohort were pooled prior to analysis in order to increase the depth coverage of the proteome and to minimize the impact of individual subjects. It should be noted that, as a primary cell, MPPMC do not maintain indefinite viability and can be maintained for a very limited number of passages (1-2 for cells from A mice, x-y for cells from Y mice, *data not shown*), thus limiting the number of cells available from each subject. Pooled lysates, 100 μg each cohort, were concentrated and trypsin digested using the S-Trap protocol (Protifi).[22]

### Western Blots & Immunofluorescence

MC are unique among normal cells in that they display both epithelial and mesenchymal characteristics, a property usually limited to regenerating or neoplastic cells. To validate the epithelial and mesenchymal properties, isolated MPPMCs were evaluated by western blotting and immunofluorescence staining for expression vimentin (mesenchymal marker) and cytokeratin 18 (epithelial marker).[23],[24],[25] For immunofluorescence experiments, freshly isolated MPPMCs were cultured on 22mm^2^ glass coverslips (coated with rat tail collagen type I), washed twice with ice-cold PBS, and fixed with 4% paraformaldehyde in 0.12M sucrose in PBS for 20 minutes at room temperature. Cells were blocked with 3% Bovine Serum Albumin (BSA) in PBS for 1 hour at room temperature, incubated with primary antibody (1:100) in 3% BSA for 1h at 37°C, rinsed thrice for 5 minutes with PBS, and incubated with appropriate Alexa-Fluor conjugated secondary antibody at a 1:300 dilution for 30 minutes at 37°C. After washing, cells were allowed to dry, mounted with VECTASHIELD Mounting Media with DAPI (Vector laboratories), and visualized on a Leica DM5500 B Fluorescence Microscope.[26] Images presented here were cropped from the full visual field. Brightness and contrast were adjusted equally across all images.

For western blots, protein concentration of cells lysed with mRIPA buffer containing protease inhibitor was measured using DC™ Protein Assay (Bio-Rad). Protein (20μg) was electrophoresed on 9% SDS-polyacrylamide gels and transferred to methanol-activated polyvinylidene membranes. After transfer, membranes were blocked with 5% milk in TBST (150mM NaCl, 25mM Tris, 0.05% Tween 20 for 1 hour at room temperature) to prevent non-specific binding. Primary antibodies, vimentin (Sigma-Aldrich, catalog #: V5255, 1:1000) and cytokeratin 18 (Abcam, catalog #: ab668, 1:500) were diluted as indicated in 5% milk/TBST and incubated overnight at 4°C. After washing, the membranes were incubated with horseradish peroxidase-conjugated secondary antibodies (1:4000 dilution) for 1 hour at room temperature, and then visualized with chemiluminescence using ImageQuant™ LAS4000.[26],[27] Gels presented here (**Figure 2**) were equally adjusted for brightness and contrast and uncropped gel lanes are available in SI. Grouping of gels was performed, they are delineated with spacing as shown in **Figure 2**.

**Figure 2.**
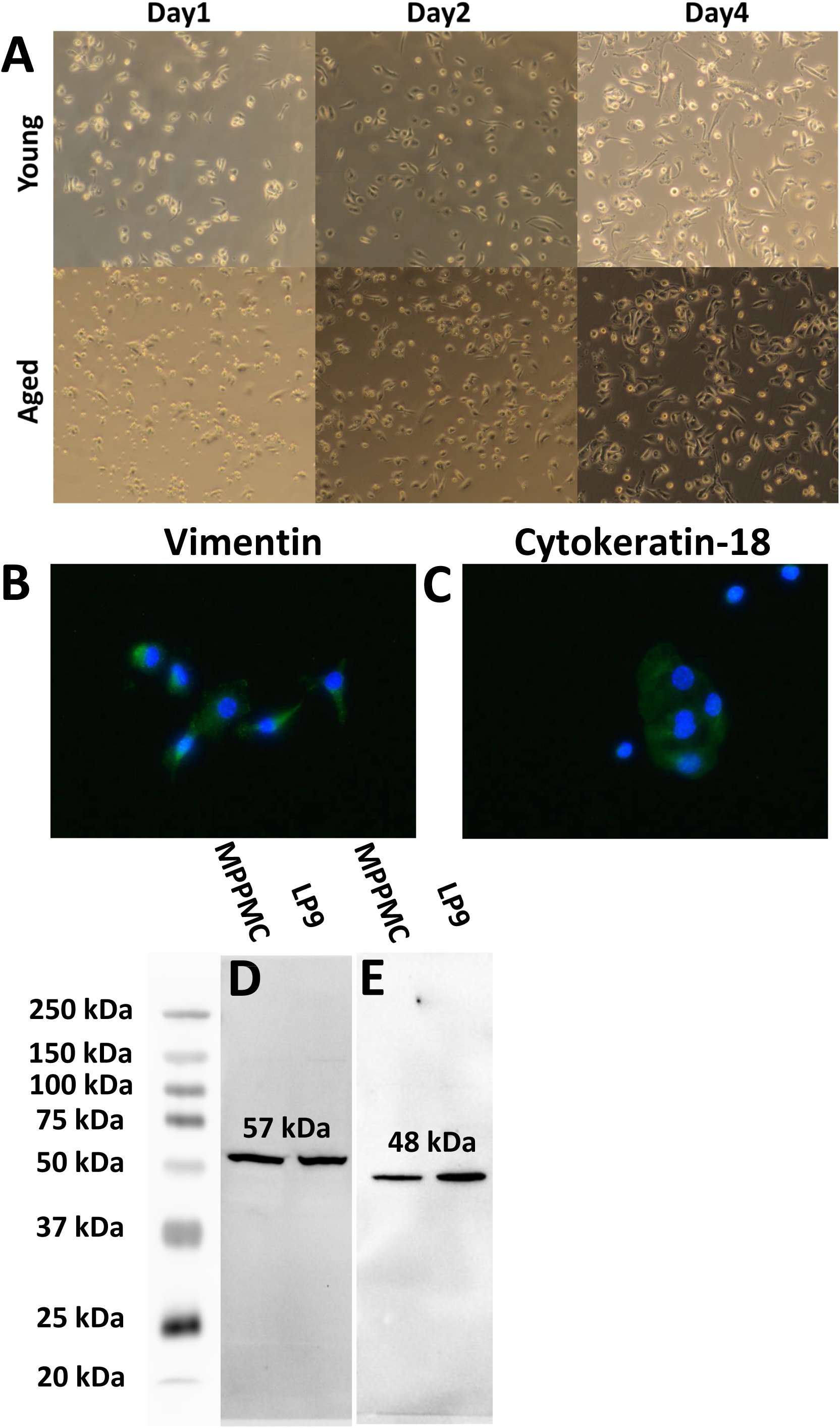
Validation of murine primary peritoneal mesothelial cells (MPPMC). **(A)** Representative morphology of MPPMC cultures collected from young (top panels) or aged (lower panels) mice shown at day 1, 2, and 4, as indicated (100X magnification). **(B)** Immuno-fluorescent staining of MPPMCs using anti-vimentin antibody (1:100 dilution) in 3% BSA for 1h at 37°C, followed by Alexa-Fluor conjugated secondary antibody (1:300) for 30 minutes at 37°C. Slides were mounted with VECTASHIELD Mounting Media with DAPI. **(C)** Immunofluorescent staining of MPPMC using anti-cytokeratin-18 antibody (1:100 dilution) in 3% BSA for 1h at 37°C, followed by processing as in (B). **(D&E)** Western blots of MPPMC and LP9 (human mesothelial cell line) cell lysates (10 ug protein). Lysates were electrophoresed on 9% SDS-PAGE and electroblotted to Immobilon membranes. After blocking with 5% milk/TBST, blots were probed with antibodies directed against (D) vimentin (1:1000 dilution) or (E) cytokeratin-18 (1:500 dilution), washed and incubated with horseradish peroxidase-conjugated secondary antibodies (1:4000) for 1 hour at room temperature. Blots were developed using Super Signal West Dura Extended Duration Substrate (Thermo) and visualized using ImageQuant™ LAS4000.

### MPPMC Proteomic Analysis

For each cohort, pooled lysates were denatured, reduced and alkylated with iodoacetamide and digested using S-Traps following manufacturers recommendations. [22] Samples were analyzed in technical triplicate per pooled cohort for a total of 12 analyses.

Digested samples were analyzed at the University of Notre Dame Mass Spectrometry core facility. Bottom-up proteomics experiments were conducted with a Thermo-Finnegan Q-Exactive (QEHF) mass spectrometer coupled to a Waters MClass ultrahigh pressure liquid chromatography system via a nanoelectrospray ionization source. Tryptic peptides were harvested from S-Trap columns and the solvent was removed by evaporation. The peptides were re-dissolved in solvent A (0.1% formic acid in water, Burdick & Jackson, MI). The volume was adjusted according to the original protein content to normalize the sample concentration. A sample containing tryptic peptides (1 µg/ul) was eluted from an Acquity BEH C18 column, 1.7-μm particle size, 300 Å (Waters) column (100 μm inner diameter × 100 mm long) using a 100-min gradient at a flow rate of 0.9 μL/min [4–33% organic solvent B (0.1% formic acid in acetonitrile), Burdick & Jackson] for 90 min, 33–80% B for 2 min, constant at 80% B for 6 min, and then 80–0% B for 2 min to re-equilibrate the column. Data were collected in positive ionization mode. Mass spectra were acquired in the Orbitrap using a TOP17 Data Dependent Acquisition (DDA) method with 60k resolving power and tandem mass spectra were then generated for the top seventeen most abundant ions with charge states z = 2-5 inclusive. Fragmentation of selected peptide ions was achieved via higher energy collisional dissociation (HCD) at normalized collision energy of 28 eV in the HCD cell of the QEHF.

### Database Analysis

PEAKS Online software (Bioinformatics Solutions Inc.) filtered to a 1% false discovery rate (FDR) was used to identify proteins present in each sample by matching tandem mass spectra with peptides expected for proteins from the Uniprot mus (mouse) database and common contaminants (Approximately 56,000 entries). Fixed modification of Carbamidomethylation (C) and variable modification of deamiadation (NQ) phosphorylation (STY), oxidation (M) and pyroglutamic formation (QE) were considered as possible post-translational modifications. We utilized the PEAKS Label Free Quantitation to make pairwise comparisons, “Female Young vs. Female Aged”, “Female Young vs. Male Young”, “Male Aged vs. Female Aged”, and “Male Young vs. Male Aged.” From each comparison, proteins differentially expressed (>2-fold) with high significance (ANOVA, -log_10_ *P value* > 10^-1.3^, 95% CI) were selected. Venn diagrams were generated using the online tools from UGent Bioinformatics & Evolutionary Genomics.[28] Gene ontology enrichment analysis was performed on those proteins for their biological processes using the ShinyGO online software (FDR cutoff α=0.05, Pathway size 2-2000).[29]

## 3. RESULTS AND DISCUSSION

### Primary Cell Culture

This study presents a reproducible method for isolation and short-term primary culture of peritoneal MC from young and aged mice [**Figure 2A**]. A unique aspect of MC is that they display both epithelial and mesenchymal characteristics. Analysis of MPPMC using immunocytochemistry shows that all DAPI-positive nuclei are associated with positive staining for cytokeratin 18 and vimentin [**Figure 2B & 2C**], indicating the lack of potentially contaminating macrophages or fibroblasts (cytokeratin 18 negative).[24] Expression was confirmed by western blotting [**Figure 2D & 2E**]].[23],[24],[25] We observed that MPPMC from young mice generally grow and divide faster than those from old mice. Our results also show that young primary cells survive longer relative to cells obtained from old animals (> 1 month *vs* ∼3 weeks, respectively, *data not shown*). Little detachment, cell death, or crowding of the plate was observed in the first 3 weeks in culture. These primary cell cultures represent an *in vitro* model to enable examination of the impact of host sex and/or age on the interaction between the mesothelium and foreign materials such as asbestos fibers or other cell types including bacteria and metastatic cancer cells with less confounding factors relative to *in vivo* experiments.

### Proteomics

MPPMC lysates from unpassaged day 4 primary cultures were concentrated and trypsin digested using S-Traps. Bottom-up proteomic analyses were then performed as described in order to compare expressed proteins in the four experimental groups: Female Young (FY), Female Aged (FA), Male Young (MY) and Male Aged (MA). Each cohort, FA, FY, MA, MY, contains a total of 2721, 1982, 2484, 2364 proteins and 899, 662, 816, 805 protein groups respectively. FA, FY, MA, MY, have 379, 58, 202, 423 proteins and 150, 17, 66, 137 protein groups [**Figure 3 & Table 1**] unique to their individual group, respectively. This represents proteins identified with at least one peptide-sequence-match with a false discovery rate of 1%. Detailed protein information is found in **Supplemental Tables 1-4** and has been uploaded to MassIVE as indicated in data availability. Gene Ontology enrichment analysis was performed separately on the highly significant proteins as determined by ANOVA in PEAKS, filtered to include those unique to each group [**Figure 4A-D**]. In the males, a majority of proteins specific to MA are involved in protein/large molecule transportation and localization. The rest of MA and a majority of MY proteins are involved in translation and a variety of metabolic processes. On the other hand, the proteins specific to FA are mostly involved in multiple metabolic pathways. The number of unique proteins in FY is too small for informative analysis.

**Figure 3.**
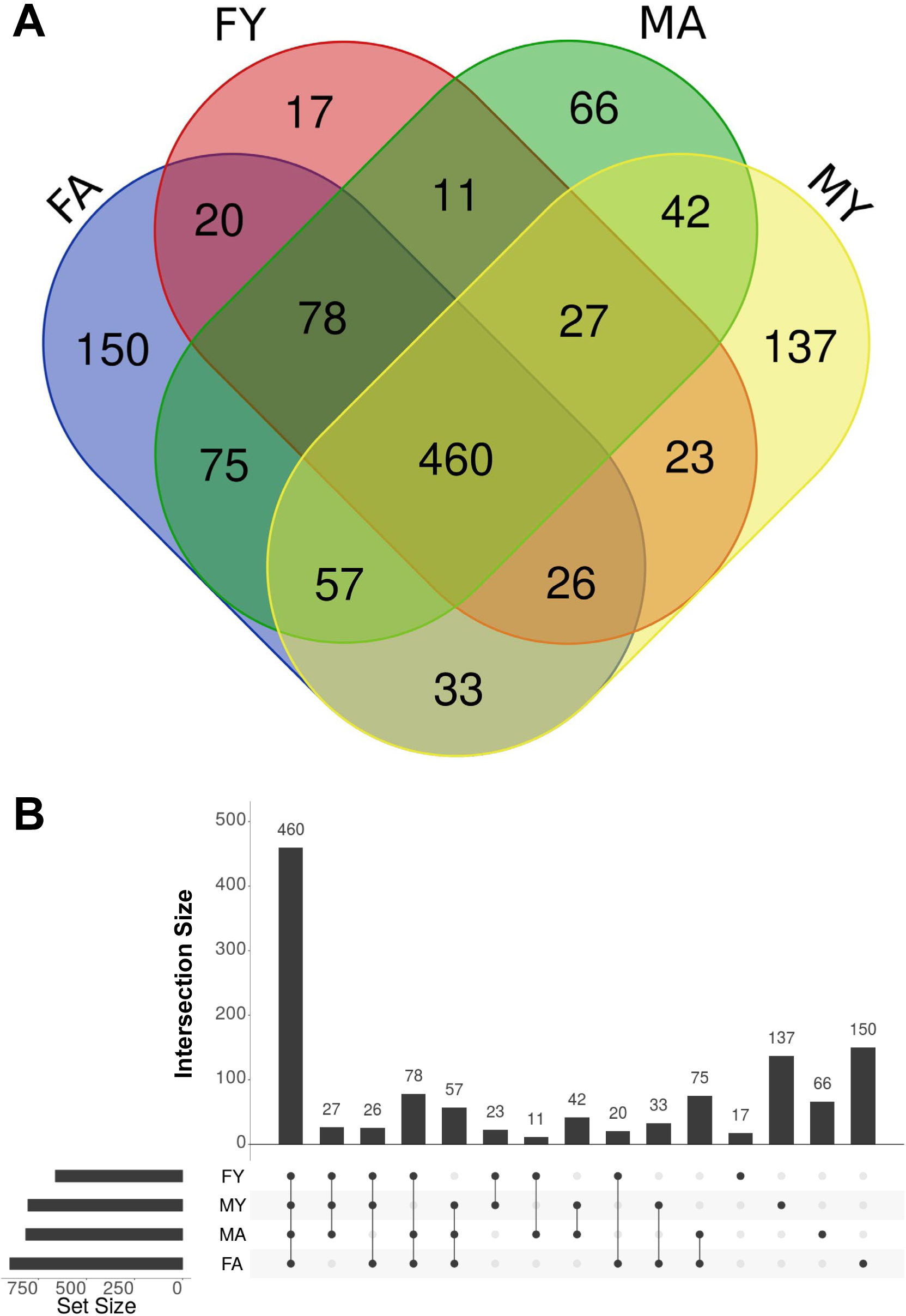
Overview of proteomic data. **(A)** A Venn diagram showing common/unique protein groups between cohorts FA, FY, MA, and MY, having 150, 17, 66, 137 protein groups unique to their individual group, respectively. **(B)** An upset plot (UpSetR Shiny App) comparing the protein groups identified from the 4 cohorts (49).

**Figure 4.**
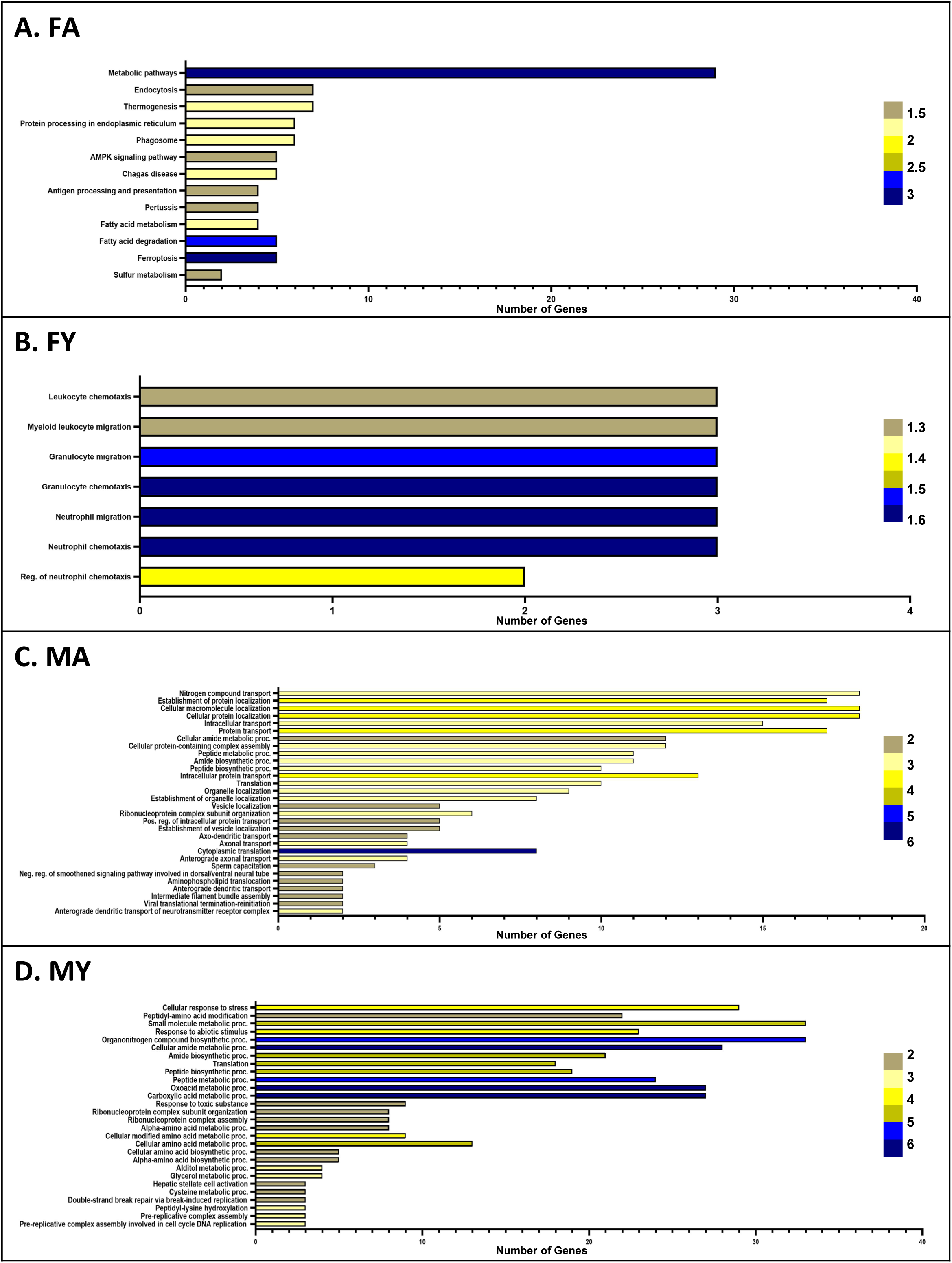
Gene Ontology Enrichment (GOE) analysis on proteins exclusively to each cohort showing their corresponding biological processes. Results show histograms listing biological processes that proteins unique to **(A)** FA, aged female; **(B)** FY, young female; (C) MA, aged male; **(D)** MY, young male; are involved. The number of genes involved in each process is shown on the x-axis. Fold enrichment increases from top to bottom. –log_10_(FDR) increases from yellow to blue.

**Table 1.** MS Results of the 4 Cohorts. Samples from each cohort underwent three replicate injections. A total of 3,697 proteins and 1,215 protein groups were identified. Each cohort, FA, FY, MA, MY, contains 2,721, 1,982, 2,484, 2,364 proteins and 899, 662, 816, and 805 protein groups respectively.

| Sample Name | Run | MS1 | MS2 | PSM | Peptides | Proteins | Protein Groups |
| --- | --- | --- | --- | --- | --- | --- | --- |
| All (4 samples) | 12 | 189219 | 184794 | 24531 | 5137 | 3697 | 1215 |
| Female Aged (FA) | 3 | 45878 | 49256 | 7278 | 3256 | 2721 | 899 |
| Female Young (FY) | 3 | 48673 | 42087 | 4681 | 2162 | 1982 | 662 |
| Male Aged (MA) | 3 | 45286 | 50454 | 7330 | 3233 | 2484 | 816 |
| Male Young (MY) | 3 | 49400 | 42997 | 5242 | 2594 | 2364 | 805 |

For each pairwise comparison (primary comparisons), the PEAKS Label Free Quantitation uses one of the groups as the “base” and measures the fold change and significance (-10log_10_(*p-value*)) of each protein from another group as compared to the “base”. The results are collected in Venn diagrams and volcano plots [**Figure 5A-D]** representing proteins up- (blue) or down- (yellow) regulated as compared to the “base”.

**Figure 5.**
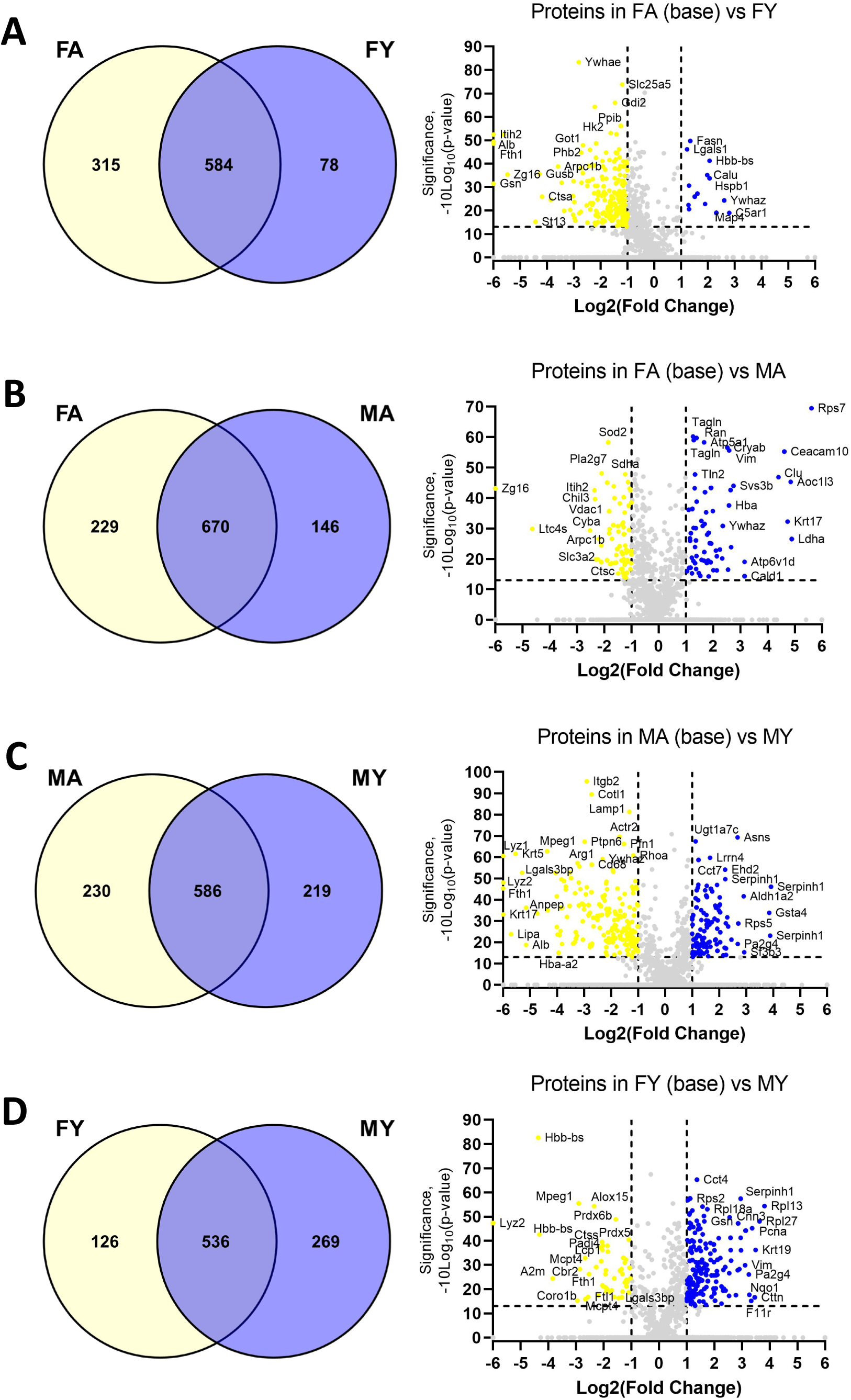
Pairwise comparisons between gender- and age-cohorts. **(A)** FA vs. FY, **(B)** FA vs. MA, **(C)** MA vs. MY, **(D)** FY vs. MY. **(Left Panels)** Common and unique proteins are shown as Venn diagrams. **(Right Panels)** Volcano plots present proteins significantly (-10log_10_(p-value) > 13) up-/down-regulated (log_2_(fold change) > 1). Proteins up-regulated when compared to the “base” are in blue, while those down-regulated are in yellow. The gene names for proteins with high significance and fold change are annotated.

All proteins having 64-fold-change or more, including ‘infinity’, are binned on the left and right margins of each plot. In each of the primary pairwise comparisons, the list of proteins with high fold-change and significance (blue and yellow) are extracted. Gene Ontology enrichment analysis showing the biological processes prevalent in each list of proteins was also performed [**Figure 6**]. The 4 lists of proteins/biological processes generated from primary comparisons were then subjected to 2 secondary comparisons. In comparison A [**Figure 6A**], “FA vs. MA”, <u>gender specific differences in aged mice</u>, was compared to “FY vs. MY”, <u>gender specific differences in young mice</u>. Their common biological processes indicate <u>gender specific differences in all ages</u>. In comparison B [**Figure 6B**], “FA vs. FY”, <u>aging specific differences in female mice</u>, is compared to “MA vs. MY”, <u>aging specific differences in male mice</u>. Their common biological processes indicate <u>aging specific differences in all sexes</u>. Comparison A shows that male and female MPPMCs are different in their protein translation and peptide metabolism pathways. Comparison B indicates that the aged and young mesothelial cells have distinct metabolic processes. Interestingly, such results have a good correlation to the gene ontology enrichment analysis on the unique proteins in each cohort [**Figure 4**].

**Figure 6.**
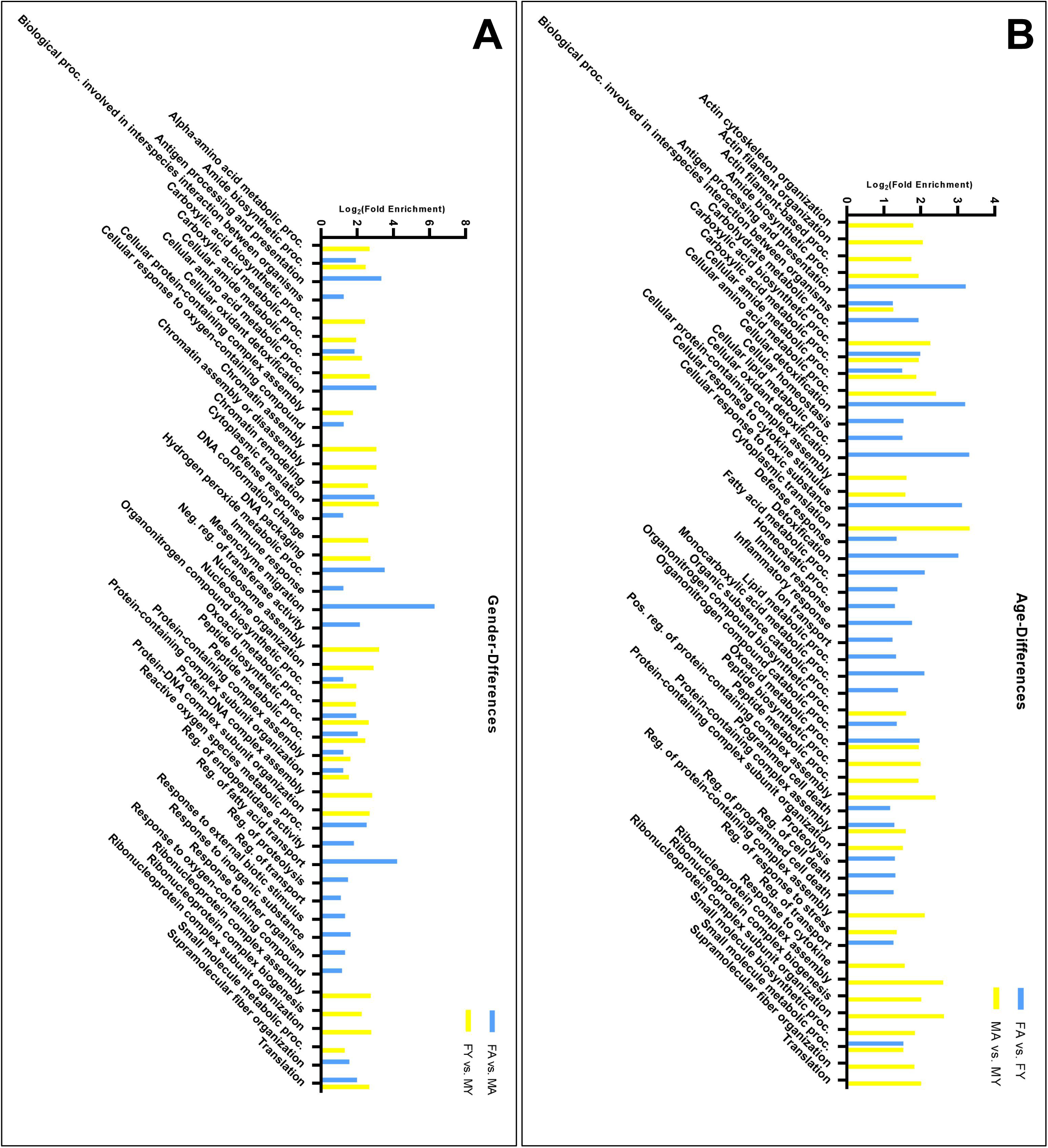
Secondary comparisons between the pairwise (primary) comparisons from Figure 5. Comparisons use only proteins with high significance (-10log_10_(*p-value*) > 13) and fold change (log_2_(fold change) > 1) as shown in Figure 5. Gene Ontology Enrichment analysis was performed on each of the 4 primary comparisons. The results are shown as histograms. Each character on the x-axis represents one biological process. The y-axis indicates log_2_(Fold Enrichment). **(A)** Comparison ⍰, FA vs. MA (blue) is compared to FY vs. MY (yellow); **(B)** Comparison ②, FA vs. FY (blue) is compared to MA vs. MY(yellow). Any character that has both a blue and yellow bin represents a common biological process shared by the 2 groups.

Our results identified differentially expressed proteins previously associated with peritoneal diseases including YWHAZ (tyrosine 3-monooxygenase/tryptophan 5-monooxygenase activation protein zeta), GSN (gelsolin), VIM (vimentin), and LGALS3BP (galectin 3 binding protein). It has been demonstrated that mesothelial cells treated with the secretome from malignant mesothelioma cultures decrease expression of YWHAZ and increase LGALS3BP levels.[30] In another study, YWHAZ, GSN, VIM were enriched in the mesothelium in a rat model of chronic peritoneal dialysis.[31] YWHAZ affects many vital cellular processes, including but not limited to metabolism, signaling, apoptosis, and cell cycle regulation.[32] Newly characterized miRNAs, miR-1-3P and miR-22, have been identified to target YWHAZ and inhibit metastasis of colorectal cancer and hepatocellular carcinoma respectively.[33],[34] YWHAZ also modulates glycolysis and promotes ovarian cancer metastasis.[32] YWHAZ is up regulated in FY and MA when both compared to FA and in MA compared to MY. Future studies may address the role of YWHAZ in cancer peritoneal metastasis, particularly in the MA host.

GSN is a multifunctional actin-binding protein and a substrate for extracellular matrix modulating enzymes.[35],[36] Outside the cell, GSN also play a role in the presentation of lysophosphatidic acid and other inflammatory mediators to their receptors. [35],[36] Many patients with renal diseases require constant automated peritoneal dialysis. It was discovered that their effluent after each peritoneal dialysis have simultaneous decrease in GSN levels, representing the presence of chronic inflammation. GSN interacts with MMP14 to enhance the activation of MMP2, thereby promoting the invasion and metastasis of hepatocellular carcinoma.[37] In contrast, GSN is also shown to have a tumor invasion suppressor role in colon cancer.[38] In our study, both FA and MY express more GSN than FY.

VIM is a mesenchymal protein and widely accepted as a biomarker for epithelial– mesenchymal transition.[39] It maintains cytoskeleton organization and focal adhesion stability.[40],[41] VIM is up regulated in MA and MY compared to FA and FY respectively, a clear gender difference that may account for many gender-specific phenotypes in cancer peritoneal metastasis. LGALS3BP is down regulated in MY when compared to MA and FY. LGALS3BP is a large oligomeric protein originally identified as a tumor-secreted antigen[42],[43] associated with inflammatory processes.[44] The majority of LGALS3BP proteins are heavily glycosylated and secreted to interact with the extracellular matrix.[45] It is associated with an IFN-induced signaling scaffold during viral infection, as well as certain bacterial proteins within infected cells.[45],[46],[47] Low expression of LGALS3BP implicates malignant progression and poor prognosis of colorectal cancer patients.[48] Similar results have been found in Ewing’s sarcoma. Patients with tumors expressing high levels of LGALS3BP display a lower risk of developing metastasis and dying.[42] Cancer cells overexpressing LGALS3BP were unable to form metastasis when injected into mice.[42]

## 4. CONCLUDING REMARKS

To our knowledge, this represents the first sex- and age-based comparison of the primary murine mesothelial cell proteome. The study provides valuable baseline information for a wide range of future studies on the role and function of peritoneal mesothelial cells in health and disease at two important physiologic age points. Future studies could compare the current data using healthy MPPMC to cells obtained from disease-bearing models to facilitate the understanding of key mesothelial cell components in peritonitis, mesothelioma, and cancer intra-peritoneal metastasis. Importantly, segregation of the data by both sex and age could reveal novel contributory factors to specific disease states involving the peritoneal cavity.

A limitation of this study design is the lack of extracellular matrix and secreted proteins in the samples analyzed. In addition, the mesothelial microenvironment *in vivo* is not only comprised solely of mesothelial cells. The collagen coating on the culture dishes used for plating of primary cell cultures does not completely mimic the more complex collagen matrix in live animals. Moreover, we recently demonstrated changes in sub-mesothelial collagen ultrastructure in aged mice relative to young using second harmonic generation and scanning electron microscopy.[21] Fibroblasts and immune cells are also not represented in the primary culture system, but could impact the mesothelial cell proteome *in vivo* due to signaling induced by soluble factors or cell:cell interactions. Future studies incorporating other aspects of the *in vivo* mesothelial cell microenvironment will generate a more complete portrait of the mesothelial proteome.

## 5. ASSOCIATED DATA

Raw and processed data are available through the MassIVE data exchange and cross-posted to ProteomeExchange. MSV000092134 ftp://MSV000092134@massive.ucsd.edu (For Review) Password: MouseAge2302* https://massive.ucsd.edu/ProteoSAFe/static/massive.jsp

## ACKNOWLEDGEMENTS

This study was supported in part by the Patterson Endowment for Excellence (Z.W.), the Walther Cancer Foundation Cancer Cure Ventures grant (R.S.), research grants RO1CA109545 and UO1CA236979 from the National Institutes of Health/National Cancer Institute (M.S.S.), and the Samuel Waxman Cancer Research Foundation (M.S.S.). Experiments were performed in the University of Notre Dame Mass Spectrometry and Proteomics Facility. Figure 1 was created with BioRender.com

## LIST OF ABBREVIATIONS

MC: mesothelial cells
FA: aged female
FY: young female
MA: aged male
MY: young male
GI: Gastrointestinal
MPPMC: murine primary peritoneal mesothelial cells
IACUC: Institutional Animal Care and Use Committee
i.p.: intra-peritoneally
DMEM: Dulbecco’s Modification of Eagle’s Medium
Pen/Strep: penicillin streptomycin solution
EGF: Epidermal growth factor
QEHF: Thermo-Finnegan Q-Exactive High Field
HCD: higher energy collisional dissociation
YWHAZ: tyrosine 3-monooxygenase/tryptophan 5-monooxygenase activation protein zeta
GSN: gelsolin
VIM: vimentin
LGALS3BP: galectin 3 binding protein.

